# Extent of similarity between agricultural and natural land covers shapes how biodiversity responds to agricultural expansion at landscape scales

**DOI:** 10.1101/2020.04.22.055111

**Authors:** Scott Wilson, Niloofar Alavi-Shoushtari, Darren Pouliot, Gregory W. Mitchell

## Abstract

The impact of agriculture on biodiversity depends on the extent and types of agriculture and the degree to which agricultural land contrasts with the natural ecosystem. Most research on the latter comes from studies on the influence of different agricultural types within a single ecosystem with far less study on how the natural ecosystem context shapes the response of biodiversity to agricultural production. We used citizen science data from agricultural areas in Canada’s Eastern Hardwood-Boreal (forest ecosystem, n=108 landscapes) and Prairie Pothole (prairie ecosystem, n=99) regions to examine how ecosystem context shapes the response of avian species diversity, functional diversity and abundance to the amount of arable crop and pastoral agriculture at landscape scales. Avian surveys were conducted along 8km transects of Breeding Bird Survey routes with land cover assembled within a 20km^2^ landscape around each transect. The amount of agriculture at which species diversity peaked differed between the forest (15%) and prairie (51%) ecosystems, indicating that fewer species tolerated the expansion of agriculture in the former. In both ecosystems, functional diversity initially increased with agriculture and peaked at higher amounts (forest: 42%, prairie: 77%) than species diversity suggesting that functional redundancy was lost first as agriculture increased. Species turnover with increasing agriculture was primarily among functional groups in forest where a shift from a low to a high agriculture landscape led to a decline in the percent of the community represented by Neotropical migrants, insectivores, upper foliage gleaners and bark foragers, and an increase in the percent of the community represented by short-distance migrants, granivores, omnivores and ground gleaners. There were few distinct shifts in the percent of the community represented by different functional groups in the prairie ecosystem. Total abundance was the least sensitive measure examined in both ecosystems and indicated that species losses with agriculture are likely followed by numerical compensation from agriculture tolerant species. Our results highlight the importance of ecosystem context for understanding how biodiversity is affected by agricultural production with declines in diversity occurring at lower agricultural extents in ecosystems with lower similarity between natural and agricultural land covers. These findings allow for more specific conservation recommendations including managing for species intolerant to agriculture in prairie ecosystems and limiting the expansion of high contrast agriculture and the loss of semi-natural habitat, such as hedge rows, in historically forested ecosystems.

---

Agricultural expansion is a key driver of global biodiversity loss (Donald et al., 2001; Kremen et al., 2002; Tscharntke et al., 2005) and these impacts are expected to increase as the human population reaches an estimated 9 billion by mid-century (Godfray et al. 2010). The manner by which agriculture impacts biodiversity can be described along two axes of influence (Cunningham et al. 2013). Along one axis is the extent to which agricultural land replaces natural land covers (‘extensification’ axis). The negative impacts of extensification across taxa are well recognized globally and include effects such as declines in species diversity and/or abundance (Tilman 1999, Donald et al. 2001), community homogenization (Ekroos et al. 2010, Karp et al. 2012, Endenburg et al. 2019) and the loss of ecosystem services (Kremen et al. 2002). Along the other axis is the similarity of the agricultural production system to the natural ecosystem into which agriculture expanded (‘similarity’ axis). Similarity in this context refers to the difference in the vertical and horizontal distribution of vegetation between the production system and the natural ecosystem and can influence the degree to which the production system supports the original biodiversity present (Perfecto et al. 1996, Benton et al. 2003). Understanding where a particular agroecosystem lies along the extensification and similarity axes has important implications for the development of management strategies to minimize biodiversity loss as agriculture expands (Cunningham et al. 2013).

Nearly all research on how the similarity of natural and agricultural land covers affects biodiversity is examined at finer spatial scales by comparing different agricultural practices within a single regional or local ecosystem. For example, by retaining much of the overstory forest structure, shade coffee production maintains a higher fraction of tropical forest biodiversity compared to open sun coffee plantations with few to no trees (Perfecto et al. 1996, Philpott et al. 2008, Caudill et al. 2015). In regions where arable cropping occurs within naturally forested landscapes, hedge rows and woody riparian strips can help support the diversity of species that select wooded or partially wooded habitats (Bátary et al. 2010, Morandin and Kremen 2013, Wilson et al. 2017). In prairie ecosystems, small strips of native grass between arable crop fields help increase the abundance of some grassland species that are intolerant of agriculture (Evans et al. 2014). A less well studied component of the similarity axis is how the response of faunal communities to agriculture differs among ecosystems that vary in the natural vegetative communities that existed before agriculture, hereafter the ecosystem context. For example, how does the relationship between the extent of cereal crop agriculture and species diversity differ when cereal crops are grown in a forested versus a prairie ecosystem? Different relationships may arise between ecosystems because of the evolutionary history of species with the structure and composition of the land covers to which they are adapted (Filloy et al. 2010, Cunningham et al. 2013). In particular, we might expect stronger negative effects of agriculture when the contrast with the natural land covers increase because the response of species depends both on 1) the degree to which species tolerate and use the agricultural cover types and 2) the degree to which species are impacted by the structural change in the broader landscape following agricultural expansion.

In this study, we used citizen science data to examine how landscape-scale avian biodiversity responds to the expansion of agriculture depending on whether the natural ecosystem context is primarily forest versus prairie. We tested this in two ecosystems in Canada, the Eastern Hardwood-Boreal region (forest ecosystem) and the Prairie Pothole region (prairie ecosystem) (Figure 1). Agricultural types are similar in both ecosystems and primarily include arable crops such as cereal grains, corn and soy as well as pasture; thus, agricultural and natural vegetation types are structurally more similar in a prairie than a forest ecosystem.

**Figure 1.**
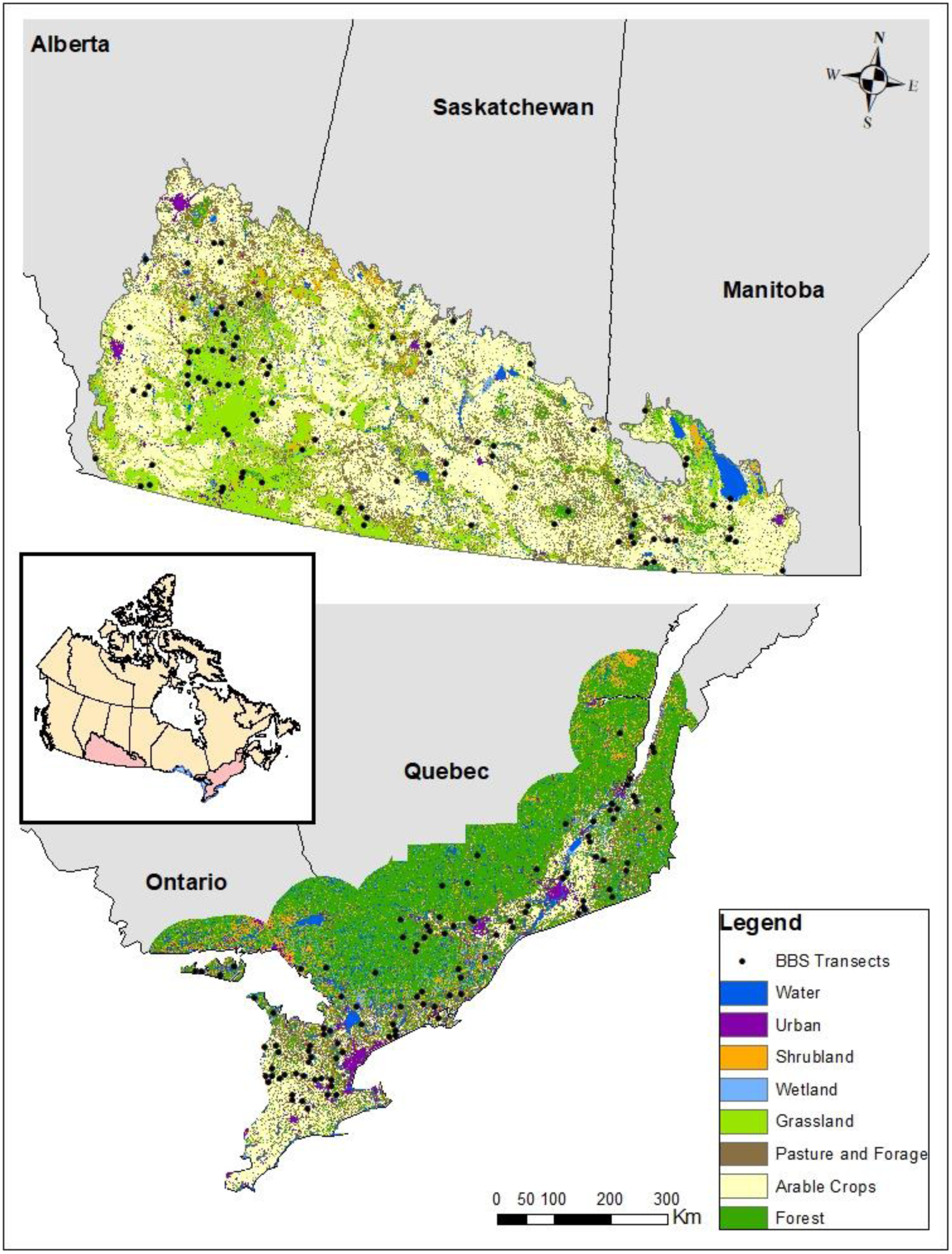
Study region showing land cover and survey transect locations in the Prairie (top) and Eastern Forest (bottom) ecosystems in western and eastern Canada respectively. The scale is the same in both panels.

Our work was based around three main questions. First, how does species diversity vary in relation to the amount of agriculture in the landscape in the two ecosystem types? Species diversity in both ecosystems might initially increase with agriculture by increasing land cover heterogeneity with few losses of species intolerant to agriculture (Bennett et al. 2006). However, a further expansion of agriculture may lead to a decline in species diversity beyond some peak as species intolerant to agriculture disappear. Our interest was how the two ecosystems differ in both the shape of the species diversity versus agriculture relationship and the threshold of agricultural amount at which species diversity begins to decline. Our second main question examined whether functional diversity is more or less sensitive than species diversity to increasing amounts of agriculture in each ecosystem. Functional diversity relates to the number, type and distribution of functions that organisms provide in an ecosystem (Díaz and Cabido 2001, Petchey and Gaston 2006). Different relationships between species diversity and functional diversity are possible along land-use gradients based on the degree of functional redundancy (Flynn et al. 2009, Cadotte et al. 2011). In particular, depending on the degree to which functionally redundant or functionally unique species are lost first as land-use intensity increases, functional diversity will decline more slowly or more rapidly than species diversity respectively (Flynn et al. 2009). Therefore, we examined the rates of change for functional versus species diversity in relation to the amount of agriculutre. We also examined how different functional groups associated with migratory behavior, foraging strata, diet, habitat (aquatic/terrestrial), nest site location and size responded to agricultural expansion in both ecosystems. Our third main question examined if there is numerical compensation across the agricultural gradient and whether it is similar in the two ecosystems. Numerical compensation occurs when some species increase in abundance across a land-use or other environmental gradient, thus numerically offsetting the loss of other species sensitive to disturbance (McGrady-Steed and Morin 2000, Touchton and Smith 2011). Numerical compensation has important implications for ecosystem resiliency if those species that increase in abundance can serve the same ecological functions as species that were lost (Naeem and Li 1997).

## METHODS

### Region characteristics

The two regions in our study are the largest agricultural regions in Canada (Fig. 1). In both, arable crops and pasture are the dominant agricultural types, although the predominance of specific crop types varies (Tables S1, S2). The Eastern Hardwood-Boreal covers a broad area of eastern Canada within the Mixedwood Plains, Boreal Shield and Atlantic Maritime ecoregions in southeastern Ontario and southern Quebec. Prior to European settlement, this region was primarily a combination of deciduous and mixed forests interspersed with lakes and wetlands. During the 18^th^ and 19^th^ centuries, deforestation increased for silviculture and agriculture. Forest regeneration has occurred since about the mid-20^th^ century particularly in eastern Ontario and southern Quebec although forest cover remains limited in southern Ontario (Butt et al., 2005). Among the transects in our survey, forests covered approximately 45% of the landscape while 31% was agriculture (Table S1). Pasture (e.g. hay, alfalfa), corn and soybean were the dominant types of agriculture in the forest ecosystem representing about 92% of total agriculture (Table S2). Agricultural lands in the region contain varying extents of semi-natural habitat (e.g. hedge rows, riparian strips) that enhance the diversity of forest and shrub-breeding avian groups (Wilson et al. 2017).

The Prairie Pothole region lies within the provinces of southern Saskatchewan, southeastern Alberta and southwestern Manitoba. This region represents the northern extension of the North American Great Plains eco-zone and is characterized by semi-arid to moist grasslands interspersed with wetlands, vegetated draws and blocks of aspen parkland (Savage 2004, Kissinger and Rees 2009). Crop farming and pastoralism began to expand rapidly toward the end of the 19^th^ and early 20^th^ century. Today the Prairie Potholes are the main agricultural region in Canada with more than 80% of national crop production (Kissinger and Rees 2009). Agriculture was the dominant land cover in the prairie ecosystem transects in our study averaging 61.9% with wheat, pasture and canola being the main types (Table S2). Grassland was the most abundant natural land cover class averaging 22.4% (Table S1).

### Avian Surveys

We utilized avian survey data collected along sections (hereafter ‘transects’) of routes from the North American Breeding Bird Survey (Sauer et al. 2017). The Breeding Bird Survey (BBS) was initiated in 1966 and is a volunteer-based survey conducted annually on over 4,000 roadside routes across North America. All surveys are conducted between late May and early July. BBS routes are 39.2km in length and contain 50 stops every 800m where the observer records all species detected within 400m during a 3 minute point count. The Breeding Bird Survey is primarily used for the analysis of long-term population trends but owing to the vast spatial and temporal coverage it is also used regularly for other macroecological studies on the impacts of climate, land-use change and disease on avian populations (Flather and Sauer 1996, LaDeau et al. 2007, Wilson et al. 2011). For the forest ecosystem, we initially selected 132 BBS routes, which included all routes from the Mixedwood Plains ecozone, the southern edge of the Boreal Shield ecozone between 70°W and 83°W, and the western edge of the Atlantic Maritimes ecozone in Quebec (Fig. 1). Routes were only retained from this region if they contained at least one year of data between 2010 and 2014 (Fig. 1). Eighteen routes were subsequently excluded from the very southern extent in Ontario (Ecoregion 7, Crins et al. 2009) as it shows distinct vegetation from the rest of the region used in our study and includes some avian species with very restricted ranges. This exlusion resulted in 114 routes for analysis; these were primarily distributed in the Mixedwood Plains (n=53) and Boreal Shield (n=45) ecozones with a smaller proportion in the Atlantic Maritime ecozone (n=16). For the prairie, we selected 109 BBS routes from the Prairie ecozone in southern Saskatchewan, southeastern Alberta and southwestern Manitoba (Fig. 1).

For all routes in the forest and prairie ecosystems, we used the bird survey information at the stop level to create two “transects” consisting of stops 1 to 11 and 21 to 31 when all land cover data was available around both transects. This process resulted in 220 transects in the forest and 203 transects in the prairie. Each transect of 11 stops was 8 km in length. We omitted stops 12 to 20 to reduce spatial autocorrelation between transects on the same route. Abundance data for each species was summed over the 11 stops within each transect and therefore transects represented the individual sampling units in this study. By using 11 stops over an 8km transect we obtained sufficient information to describe each ‘community’ in terms of species composition and the relative abundances of species, but apart from this the choice of transect length was arbitrary. We included all species detected on transects in this analysis including seven introduced species. For additional detail on the designation of transects and survey methodology see Wilson et al. (2017).

The two regions differed in the mean amount of agriculture per transect with the forest ecosystem having more transects with little to no agricultural coverage and the prairie having more transects with high agricultural coverage. To ensure a balanced representation of transects across the agricultural gradient in each region we classified all 220 transects in the forest and 203 transects in the prairie according to three categories: less than 33% agriculture (‘low’), 33-66% agriculture (‘moderate’) or greater than 66% agriculture (‘high’). In the forest this resulted in 123 low, 61 moderate and 36 high agriculture transects, while in the prairie this resulted in 33 low, 71 moderate and 99 high agriculture transects. For analyses, we then randomly selected 36 transects in the low and moderate categories in the forest ecosystem and 33 transects in the moderate and high categories in the prairie ecosystem to ensure a more balanced sample of transects across the gradient of agriculture in both ecosystems.

The methods of the BBS do not allow us to formally account for detectability in the estimation of species abundance but species may be missed during surveys or could differ in detectability due to factors such as openness, time of day or weather. To provide some accounting for the possibility that a species is present but missed during any survey, we used a 5- yr window to estimate average abundance per transect. The mid-point for this 5-yr window was based on the year when land cover was obtained: 2012 in the forest and 2015 in the prairie. Therefore, the average abundance per species were based on data from all surveys conducted on transects between 2010 and 2014 in the former and between 2013 and 2017 in the latter. As a further examination of our ability to detect all species we also estimated cumulative species richness; specifically, the number of transects required to detect 95% and 99% of the species pool in each ecosystem (see Supplemental Information, Figure S1). Once all transect data had been assembled as defined above we calculated the Shannon index (Shannon 1948) on each transect as a measure of species diversity and the summed average abundance of each species on each transect as a measure of total abundance.

### Functional Diversity

We assigned all species based on five categorical functional traits (migratory status, diet, foraging strata, habitat, nest location) to estimate relationships between functional diversity and agriculture. Hereafter we refer to the groups of species defined by these functional traits as functional groups. We also included body size measurements for overall size based on estimates of mass obtained in Sibley (2000). Migratory functional group designations included resident, short-distance migrant and Neotropical migrant based on descriptions of the wintering range from the Birds of North America (Rodewald 2018). Residents were species with no directional change in their breeding and winter distributions. Short-distance migrants were species where the winter distribution lies south of the breeding distribution but north of the Tropic of Cancer. Neotropical migrants were species that breed in Canada and primarily over-winter south of the Tropic of Cancer including the Caribbean (Hagan and Johnston 1992). In cases of species with both migratory and resident populations, we used the Birds of North America (Rodewald 2018) to classify them based on the migratory status of the populations breeding within our study region. For three species, the migratory designation differed between the two ecosystems with a species being a resident in one ecosystem and a short-distance migrant in the other (Table S3).

Diet and foraging strata were based on information in Elton Traits 1.0 (Wilman et al. 2014) and the Birds of North America (Rodewald 2018). Dietary functional group categories were assigned based on the majority diet in Wilman et al. (2014) and included herbivore, granivore, frugivore, nectarivore, insectivore, omnivore, carnivore and scavenger. Omnivores were those species where the majority diet was either plants or insects but also contained <50% vertebrates and/or scavenging. Foraging strata were based on the location where the majority of food (>50%) is obtained and included ground forager, lower foliage gleaner, upper foliage gleaner, bark forager, aerial forager, surface water forager, under water forager. Elton Traits 1.0 define foliage gleaners based on proportion of food obtained from the understory, mid-story and canopy layers and the majority of species in our analysis use two or more of these layers. We therefore defined lower foliage gleaners as those species where the proportion of time foraging in understory and mid-story exceeded the proportion of time in mid-story and canopy. Upper foliage gleaners were the reverse where the proportion of time foraging in mid-story and canopy exceeded the proportion of time in understory and mid-story.

We assigned species to one of two habitat classes depending on whether they have obligate associations with wetlands and/or waterbodies during the breeding period or whether they were primarily terrestrial, although many terrestrial species frequently use vegetation adjacent to wetlands. This habitat classification was done to separate species that primarily have a functional role in aquatic versus terrestrial systems. While foraging strata and habitat are often related, several aquatic obligates primarily forage on the ground adjacent to wetlands rather than at or below the water surface (e.g. geese) and several terrestrial species forage at aquatic environments (e.g. swallows).

We used the Birds of North America account (Rodewald 2018) to identify the most common nest site location for a species including: open-cup ground (less than 50cm off ground), open-cup above ground (>50 cm off ground in tree, shrub, marsh vegetation (e.g. cattails)), tree cavity and rock faces/cliffs. Species that now regularly use anthropogenic sites were assigned based on the typical natural sites used (e.g. Barn Swallows, *Hirundo rustica*). Because of their distinctive functional role as brood parasites without their own nests, we assigned Brown-headed Cowbirds (*Molothrus ater*) as a separate brood parasite category.

Functional diversity was estimated using Rao’s Quadratic entropy (*FD*_*Q*_, Botta-Dukát 2005) based on the six traits for each species (migratory status, diet, foraging strata, habitat, nest location, size). FD_Q_ estimates the sum of the functional dissimilarities of the species in the community on each transect weighted by the relative abundance of each species (Botta-Dukát 2005). FD_Q_ was calculated using the FD package in R (Laliberté and Legendre 2010, Laliberté et al. 2014). With diversity measures such as *FD*_*Q*_ it is difficult to recognize which specific functional traits vary across a land-use gradient and thus give rise to any observed changes in functional diversity. Therefore, to further examine how functional groups respond to the amount of agriculture we calculated the percentage of abundance represented by each functional group for the categorical traits along with size-weighted abundance on transects in low agriculture (<33% coverage), moderate agriculture (33-66%) and high agriculture (>66%) in each ecosystem. These categories correspond to our selection of a random set of transects sampled from each category in creating a balanced sample size across the agricultural gradient.

### Land cover

We used annual crop inventory data from Agriculture and Agri-Food Canada (Fisette et al. 2013) to summarize land cover at a 1km buffer on either side of each transect line including the ends. This estimation resulted in a 20km^2^ landscape surrounding each transect. The crop inventory is a 30 m resolution data set with detailed information on up to 66 classes of land cover data, including agricultural and natural vegetation classes. We determined the total number of 30 * 30 m pixels for each land cover class within the 1km buffer and converted these to proportions based on the total number of pixels in each landscape. While we have previously summarized land cover at larger 3km and 5km spatial scales, the results of earlier analyses examining the response of alpha and beta diversity to land cover amounts are very similar regardless of the scale used (Endenburg et al. 2019). We added the proportions of cropland (including cereals, pulses, fruits and vegetables) and pasture to determine the total proportion of agricultural land within the landscape around each transect.

### Statistical Analyses

We used general linear mixed models (R package lme4; Bates et al. 2015) to test each of our three questions. Akaike’s Information Criterion was used to compare support for different candidate models where a variable was considered influential if the addition of that variable resulted in a lower AIC relative to the same model without it (Burnham and Anderson 2002). Additional variables are likely to influence species detection or abundance on surveys and were first tested in all models to control for their effect prior to testing the effects of agriculture in each ecosystem. These included 1) BBS route as a random effect to account for sources of variance specific to each route such as location, observer skill, weather, and survey date, and 2) whether the survey was on transect 1 or 2 on each route as a fixed effect. This latter variable was included to account for potential effects of time of day, as transect 1 on each route is surveyed in early morning while transect 2 is surveyed in mid-morning. While accounting for these two variables we then tested for effects of ecosystem and agriculture on species diversity, functional diversity and abundance. These tested included models that included a linear and a quadratic relationship between proportion agriculture and the response variable as well as an interaction between ecosystem and the linear and quadratic effects of agriculture on the response variable. The interaction models in each case were used to test our predictions that the influence of agriculture differs between the forest and prairie ecosystems. Functional diversity tends to increase with the number of species and in this study the correlation between functional diversity and species richness was r = 0.61. Therefore, we included species richness as a covariate of functional diversity to identify the relationship with amount of agriculture while accounting for species number. In addition to the above modeling approaches, our examination of functional diversity included a comparison of rates of change between functional and species diversity as agriculture increased. We verified model assumptions through visual inspection of quantile plots and plots of residuals vs fitted values; these did not reveal any issues with lack of fit.

## RESULTS

### General Characteristics

The forest ecosystem contained 176 species detected across the 108 transects with an average of 52 species detected per transect (min=23, max=80). The prairie ecosystem contained 184 species detected across 99 transects with an average of 47 species detected per transect (min=21, max=82). The community was well sampled across both ecosystems with 95% of the species detected with coverage of any random selection of 69% of the transects in forest and 67% of the transects in prairie (Fig. S1). Average species and functional diversity was slightly higher in forest than prairie while abundance was higher in prairie than forest (Fig 2, Table S4). The coefficient of variation in all three measures was higher in prairie than forest and notably so for functional diversity (Table S4).

**Figure 2.**
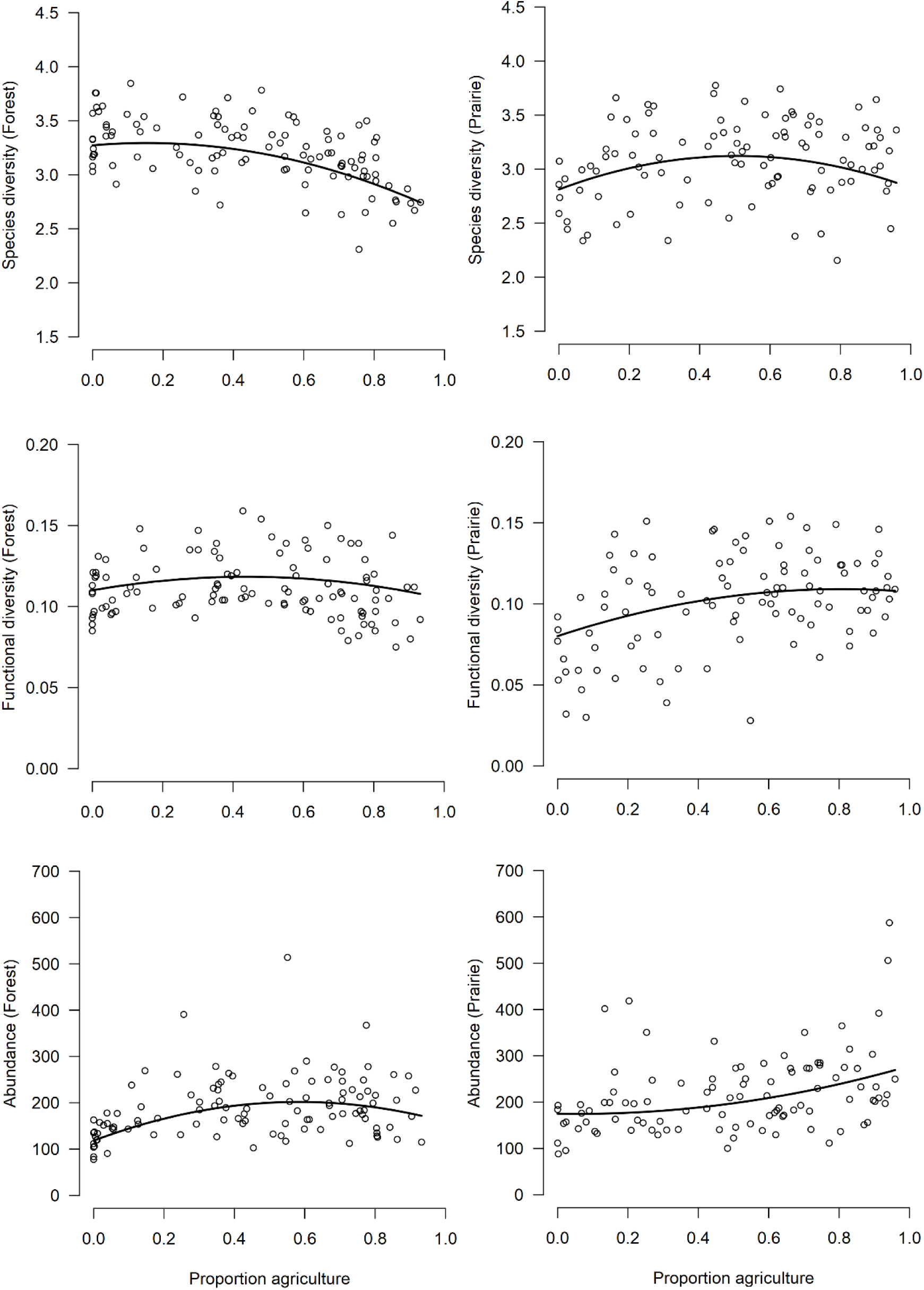
Within-landscape avian species diversity (Shannon index), functional diversity (Rao’s Quadratic entropy) and individual abundance for 20km^2^ landscapes in the Forest (n=108) and Prairie (n=99) ecosystems. The three measures were based on avian species and abundance data measured along 8km sections of Breeding Bird Survey routes within each landscape. Points are the observed estimates for each transect while the curved lines are the best fit estimates from the coefficients in the top models in Table 1.

### Species Diversity vs Extent of Agriculture

We found the strongest support for models that included an interaction between ecosystem and a quadratic response to agriculture (Table 1a). As predicted, diversity initially increased as agriculture increased in both ecosystems before declining beyond a peak (Figure 2); this peak was at a lower extent of agriculture in forest (15%) than prairie (51%). Ecosystem-specific estimates describing the relationship for forest were 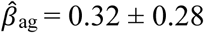 and 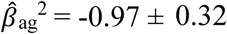, and for prairie were 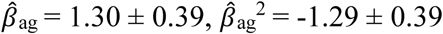. Route effects accounted for 47% of the residual variance in the top model. Time of survey was also influential with higher diversity reported on the second transect of the route 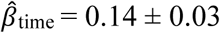.

**Table 1.** AIC model selection results for the examination of agricultural effects on species diversity, functional diversity and abundance in the forest and prairie ecosystems. k = number of model parameters, ΔAIC = change in AIC relative to top model, w_i_ = model weight.

| Variables | k | $\Delta AIC$ | $w_i$ |
| --- | --- | --- | --- |
| *a) Species Diversity* |  |  |  |
| <sup>a</sup>ecosystem*agriculture<sup>2</sup> | 8 | 0.0 | 0.99 |
| ecosystem*agriculture | 6 | 9.0 | 0.01 |
| ecosystem+agriculture<sup>2</sup> | 6 | 11.8 | 0.00 |
| ecosystem+agriculture | 5 | 15.2 | 0.00 |
| ecosystem | 4 | 16.9 | 0.00 |
| null | 3 | 17.8 | 0.00 |
| *b) Functional Diversity* |  |  |  |
| ecosystem*agriculture<sup>2</sup> | 9 | 0.0 | 0.66 |
| ecosystem*agriculture | 7 | 4.8 | 0.17 |
| ecosystem+agriculture<sup>2</sup> | 6 | 8.9 | 0.09 |
| ecosystem+agriculture | 6 | 9.6 | 0.07 |
| null | 4 | 15.2 | 0.00 |
| ecosystem | 5 | 16.6 | 0.00 |
| *c) Abundance* |  |  |  |
| ecosystem*agriculture<sup>2</sup> | 8 | 0.0 | 1.00 |
| ecosystem+agriculture<sup>2</sup> | 6 | 25.3 | 0.00 |
| ecosystem*agriculture | 6 | 27.2 | 0.00 |
| ecosystem+agriculture | 5 | 34.8 | 0.00 |
| ecosystem | 4 | 61.8 | 0.00 |
| null | 3 | 75.7 | 0.00 |

<sup>a</sup>All models included BBS route as a random variable and whether the survey was transect 1 (stops 1 to 11) or 2 (stops 21 to 31) to account for time of day. The “null” model included route and time of day without ecosystem or agriculture. Functional diversity analyses also included species richness as a covariate.

### Response of Functional Diversity vs Species Diversity

The model with the strongest support for functional diversity again included an interaction between ecosystem and a quadratic response to agriculture (Fig. 2, Table 1b), however, in both ecosystems the peak occurred at a greater extent of agriculture compared to species diversity. In the forest, functional diversity peaked at approximately 42% agriculture 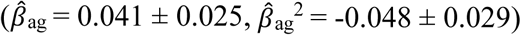, [ineq, while in prairie the peak was at 77% agriculture 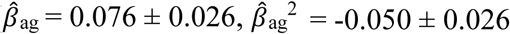, Fig 2). Route effects accounted for 37% of the residual variance. Estimates of functional diversity were higher on average on the second transect of the route 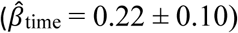. Our results indicate that in landscapes with greater natural cover than agricultural cover, functional diversity is less sensitive than species diversity to increasing agriculture. However, in the forest ecosystem, the two measures of diversity declined at nearly similar rates in landscapes with more than 80% agriculture while in the prairie ecosystem, species diversity always declined more strongly than functional diversity (Fig. 3).

**Figure 3.**
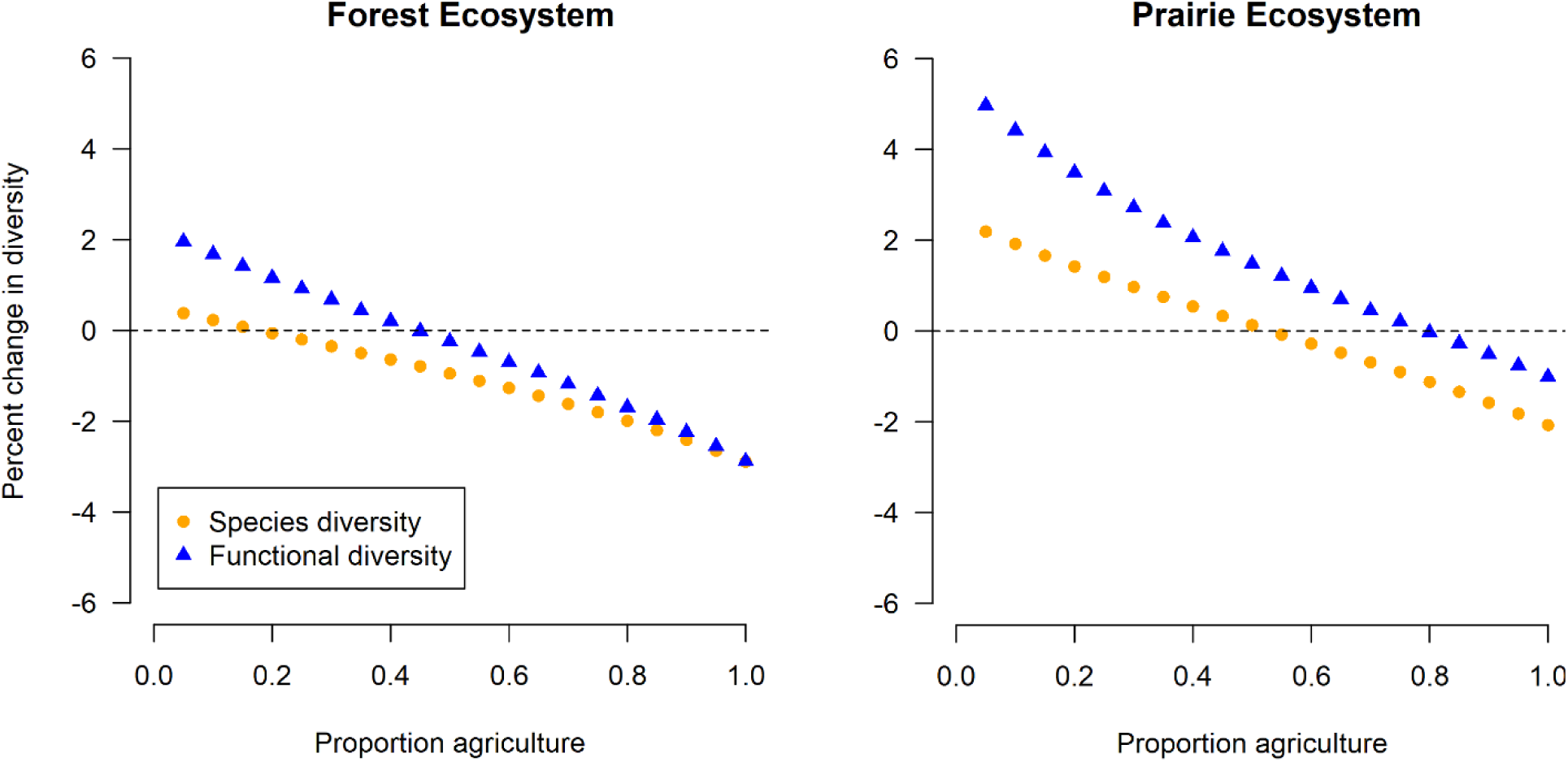
Percent change in species diversity and functional diversity for each 0.05 increase in proportion agriculture in each ecosystem. For example, the point at 0.05 is the change as agriculture increased from 0 to 0.05 cover. Values above and below the horizontal line at 0 indicate proportions of agriculture where the measures of diversity were increasing and decreasing respectively.

There were clear patterns in the susceptibility of different functional groups to agriculture in the forest ecosystem. In particular, a shift from a low to a high agriculture landscape led to a decline in the percent of the community represented by Neotropical migrants, insectivores, upper foliage gleaners and bark foragers, and an increase in the percent of the community represented by short-distance migrants, granivores, omnivores and ground gleaners (Table 2). There were few distinct shifts in the percent of the community represented by different functional groups in the prairie ecosystem. The only significant change from low to high agriculture landscapes was a decline in the proportion of the community represented by ground-nesting species and an increase in the proportion represented by open cup above ground-nesting species (Table 2).

**Table 2.** Percent of abundance by functional group in low (<33%), moderate (33-66%) and high (>66%) agriculture landscapes in the forest and prairie ecosystems. Values shown are the mean and standard error across transects within each agriculture class. Sample size of transects were equal in each class (forest, n=36; prairie, n=33). Bolded values and superscripts indicate cases where the proportion of that functional group differed among the agricultural classes based on non-overlapping 95% confidence intervals. The superscript following the estimate indicates which group it differed from (L=low, M=moderate, H=high).

| Functional group | Forest Ecosystem |  |  | Prairie Ecosystem |  |  |
| --- | --- | --- | --- | --- | --- | --- |
|  | Low | Mod. | High | Low | Mod. | High |
| *Migratory status* |  |  |  |  |  |  |
| Neotropical | **35.5 ± 2.1<sup>MH</sup>** | **21.1 ± 1.3<sup>LH</sup>** | **13.3 ± 1.0<sup>LM</sup>** | 14.2 ± 2.3 | 16.1 ± 1.6 | 18.1 ± 1.4 |
| Short-distance | **43.9 ± 1.6<sup>MH</sup>** | **52.7 ± 1.2<sup>LH</sup>** | **61.3 ± 1.6<sup>LM</sup>** | 79.1 ± 2.6 | 76.0 ± 2.0 | 72.0 ± 2.1 |
| Resident | 20.6 ± 1.3 | 26.2 ± 1.2 | 25.4 ± 1.4 | 6.8 ± 1.2 | 8.0 ± 1.1 | 9.9 ± 2.1 |
| *Diet* |  |  |  |  |  |  |
| Carnivore | 1.7 ± 0.3 | 1.6 ± 0.3 | 2.4 ± 1.0 | 2.1 ± 0.3 | 2.3 ± 0.5 | 1.8 ± 0.3 |
| Frugivore | 1.4 ± 0.2 | 1.4 ± 0.2 | 1.1 ± 0.2 | 0.3 ± 0.1 | 1.0 ± 0.5 | 0.3 ± 0.1 |
| Granivore | **17.7 ± 0.9<sup>MH</sup>** | **21.7 ± 1.0<sup>L</sup>** | **25.3 ± 1.1<sup>L</sup>** | 28.8 ± 2.1 | 25.2 ± 2.0 | 26.9 ± 1.7 |
| Herbivore | 2.5 ± 0.9 | 4.8 ± 1.2 | 4.3 ± 1.2 | 7.2 ± 1.2 | 10.4 ± 1.4 | 6.5 ± 1.9 |
|  | Low | Mod. | High | Low | Mod. | High |
| Insectivore | **58.7 ± 2.3<sup>MH</sup>** | **43.5 ± 1.6<sup>L</sup>** | **40.8 ± 1.3<sup>L</sup>** | 51.1 ± 1.7 | 47.8 ± 2.0 | 48.8 ± 1.9 |
| Nectarivore | 0.14 ± 0.04 | 0.05 ± 0.03 | 0.02 ± 0.02 | 0.02 ± 0.02 | 0 | 0.02 ± 0.01 |
| Omnivore | **17.8 ± 1.5<sup>MH</sup>** | **27.0 ± 1.6<sup>L</sup>** | **26.0 ± 1.6<sup>L</sup>** | 10.5 ± 2.1 | 13.2 ± 1.5 | 15.7 ± 1.5 |
| Scavenger | 0.06 ± 0.04 | 0.13 ± 0.06 | 0.13 ± 0.07 | 0 | 0.07 ± 0.05 | 0.03 ± 0.02 |
| *Foraging substrate* |  |  |  |  |  |  |
| Aerial | 3.3 ± 0.3 | 4.8 ± 0.4 | 4.9 ± 0.6 | 4.1 ± 0.6 | 5.5 ± 0.8 | 7.3 ± 1.3 |
| Bark | **2.8 ± 0.3<sup>MH</sup>** | **1.2 ± 0.2<sup>LH</sup>** | **0.6 ± 0.1<sup>LM</sup>** | 0.2 ± 0.1 | 0.2 ± 0.1 | 0.2 ± 0.1 |
| Ground | **51.7 ± 1.9<sup>MH</sup>** | **64.2 ± 1.1<sup>LH</sup>** | **71.9 ± 1.4<sup>LM</sup>** | 77.9 ± 2.6 | 70.2 ± 2.4 | 72.5 ± 1.7 |
| High-Foliage | **22.9 ± 1.6<sup>MH</sup>** | **9.7 ± 0.7<sup>LH</sup>** | **5.2 ± 0.5<sup>LM</sup>** | 1.9 ± 0.7 | 3.1 ± 0.9 | 1.9 ± 0.3 |
| Low-Foliage | **17.2 ± 0.8<sup>H</sup>** | **17.3 ± 0.7<sup>H</sup>** | **13.9 ± 0.6<sup>LM</sup>** | 4.9 ± 1.2 | 6.5 ± 1.0 | 8.2 ± 0.9 |
| Surface Water | **1.2 ± 0.2<sup>M</sup>** | **2.5 ± 0.4<sup>L</sup>** | 3.5 ± 1.2 | 7.6 ± 1.1 | 11.2 ± 1.4 | 8.4 ± 1.2 |
| Under Water | **0.9 ± 0.3<sup>H</sup>** | **0.4 ± 0.2<sup>H</sup>** | **0.03 ± 0.02<sup>LM</sup>** | 3.4 ± 1.0 | 3.3 ± 0.9 | 1.5 ± 0.4 |
| *Nesting location* |  |  |  |  |  |  |
| Brood Parasite | **0.3 ± 0.1<sup>H</sup>** | 0.6 ± 0.1 | **0.8 ± 0.1<sup>L</sup>** | 3.9 ± 0.5 | 3.9 ± 0.6 | 3.3 ± 0.5 |
|  | Low | Mod. | High | Low | Mod. | High |
| Tree Cavity | **11.1 ± 0.9<sup>MH</sup>** | **14.9 ± 0.9<sup>L</sup>** | **15.3 ± 1.1<sup>L</sup>** | 6.1 ± 1.1 | 7.8 ± 1.2 | 9.2 ± 1.7 |
| Ground | 33.7 ± 1.7 | 29.1 ± 1.7 | 28.8 ± 2.1 | **62.7 ± 3.4<sup>H</sup>** | **53.4 ± 2.7<sup>H</sup>** | **42.6 ± 2.0<sup>LM</sup>** |
| Open Cup | 51.5 ± 1.5 | 51.4 ± 1.4 | 50.0 ± 1.7 | **24.9 ± 2.5<sup>L</sup>** | **30.9 ± 2.3** | **38.2 ± 2.3<sup>H</sup>** |
| Rock Cavity | 3.4 ± 0.4 | 4.0 ± 0.4 | 5.1 ± 0.7 | 2.4 ± 0.5 | 4.0 ± 0.8 | 6.7 ± 1.5 |
| *Body size* |  |  |  |  |  |  |
| Mass (g) | 197.8 ± 31.6 | 285.7 ± 39.5 | 283.9 ± 38.0 | 241.0 ± 26.2 | 341.6 ± 39.7 | 252.8 ± 27.7 |

### Numerical compensation by agriculture-tolerant species

As with species diversity and functional diversity, top models included an ecosystem by agriculture interaction (Table 1c). In the forest, total abundance increased with agriculture and peaked at approximately 55% coverage before declining 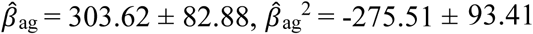, Fig. 2). The relationship was slightly increasing and near linear without a peak in the prairie 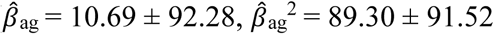, Fig. 2). Thus we found evidence for numerical compensation as agriculture increased in both ecosystems. Route effects accounted for a considerable amount (53%) of the residual variance. Abundance was higher on average on the second transect 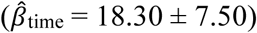.

## DISCUSSION

Studies that assess anthropogenic impacts on biodiversity at landscape-scales across different ecosystems are rare due to the effort required to survey species with a consistent methodology across broad geographic regions. We overcame this challenge by using a continental-scale citizen science program that allowed us to integrate avian biodiversity data with high resolution crop mapping data. By doing so, our study highlights the importance of the ecosystem context in the response of biodiversity to agriculture. The amount of agriculture at which biodiversity peaked and then declined occurred at higher proportions of agriculture in prairie grasslands where ecosystem similarity between agricultural and natural land covers was higher than in forests. This finding supports the idea that the effect of agriculture on community composition and diversity depends both on the degree to which species use agricultural land covers and their tolerance to land cover change. Our results also showed consistent patterns between the two ecosystems with species diversity being the most sensitive to agricultural expansion, followed by functional diversity and then abundance.

Our first main question examined whether the natural ecosystem influenced the shape of the relationship between amount of agriculture and species diversity. As predicted, species diversity initially increased with agriculture in both ecosystems. This initial increase is likely due to the fact that agriculture and associated cover types (e.g. hedge rows between fields) create additional heterogeneity without strong impacts on species that are specialists of the native ecosystem land covers. Studies on taxa in other ecosystems have emphasized the value of mosaicked landscapes to the total richness of a taxon including butterflies in Sweden (Weibull et al. 2000) and birds in Spain (Pino et al. 2000) and Australia (Haslem and Bennett 2008). Our results showed that the threshold at which the effect of agriculture on diversity changes from a benefit to a cost is lower in the forested ecosystem. This greater negative impact of agriculture on species diversity in a forest versus prairie ecosystem is likely due to at least two factors. First, habitat selection is typically a multi-scale process where initial selection is based on landscape structure followed by finer scale selection of a habitat patch within that landscape (Jones 2001, Farrell et al. 2019). A higher proportion of species in forest will be adapted to wooded environments and agriculture likely affects these species at both spatial scales. In other words, expanding agriculture initially filters out woodland species that do not tolerate the conversion to a semi-open landscape and of those species that do tolerate greater openness, only a subset use agricultural land covers or adjacent field edge habitats like hedge rows at finer scales (Endenburg et al. 2019). In contrast, the general openness of agricultural landscapes in prairie differs less from that of the native ecosystem (i.e., greater similarity) and thus the effect on species diversity may be determined primarily by the extent to which species are influenced by local tolerance to agricultural land covers rather than landscape characteristics (e.g. Koper and Schmiegelow 2006). Second, species diversity is relatively lower in natural grassland landscapes of the prairie ecosystem (i.e. landscapes with very low agriculture) than in natural forest landscapes of the Eastern Hardwood and Boreal region (see Fig. 2). This lower diversity may be related to the simpler vegetation structure of grasslands relative to forests, as less structurally diverse vegetation may provide fewer resources and support fewer species (Tews et al. 2003). With lower diversity in natural prairie landscapes, there may be more opportunity for agricultural habitats to provide additional heterogeneity that benefits a range of species and results in a higher proportion agriculture at which diversity peaks.

Our second main question examined the relationship between functional diversity and agriculture and how that relationship compared to that for species diversity. Functional diversity initially increased with agriculture in both ecosystems reaching a predicted peak at 42% agriculture in the forest ecosystem and 77% agriculture in the prairie ecosystem before declining as agriculture increased further. In both cases, the fact that functional diversity peaked at a greater proportion of agriculture than species diversity suggests that species loss was primarily by functionally redundant species (Flynn et al. 2009). The range in proportion of agriculture at which this was predicted to occur was 15-42% in forest and 51-77% in prairie (i.e. the range where species diversity declined while functional diversity was still increasing). The greater sensitivity of species diversity to agriculture was maintained in the most extensive agricultural landscapes in the prairie ecosystem but in the forest ecosystem the two measures declined similarly beyond about 80% agriculture suggesting that the redundancy in the avian community had been lost. We acknowledge an important caveat in this comparison of the two measures in that estimates of functional diversity are dependent on how species are assigned to functional groups (Mayfield et al. 2010, Cadotte et al. 2011). The patterns we observed are those given the six traits considered in this study, which were selected to reflect the range of species in the community and their role in the ecosystem throughout the annual cycle. However, these species-functional diversity patterns may have differed had we estimated functional diversity based on fewer or more traits.

The decline in functional diversity beyond the peak proportion agriculture in forest was largely related to a decline in Neotropical migrant, foliage-gleaning insectivores resulting in a community with greater proportional representation from short-distance migrants, granivores, omnivores and ground gleaners. In contrast, there were few clear patterns for the response of functional groups to agriculture in the prairie ecosystem suggesting that changes in species diversity across gradients of agriculture were likely due to a turnover of species abundance within rather than among functional groups as observed in the forest ecosystem. As an example, within the ground-gleaning insectivores and granivores in the prairie ecosystem, tolerance to agriculture varies widely ranging from species intolerant to cropland, such as Chestnut-collared Longspur (*Calcarius ornatus*)(Bleho et al. 2015) to more tolerant species, such as Vesper Sparrow (*Pooecetes gramineus*)(Jones and Cornely 2002).

Our third main question examined the potential for numerical compensation in each ecosystem whereby more tolerant species increase in abundance to offset declines in the abundance of less tolerant species. We found evidence for numerical compensation in both ecosystems. Abundance peaked at 55% agriculture in the forest ecosystem and declined thereafter in the most extensive agricultural landscapes. In the prairie ecosystem, abundance continued to increase with increasing agriculture despite losses in species and functional diversity when agriculture covered 75-100% of the landscape. However, the process by which species compensated numerically differed between the two. In the forest ecosystem it was largely due to a turnover in the functional groups representing the community as described earlier. Only a small group of species were among the most abundant species in the community in both natural and extensive agricultural landscapes. In contrast, compensation in the prairie ecosystem was largely due to an increase in abundance from species that also readily used natural landscapes (e.g. Red-winged Blackbird (*Agelaius phoenicus*), Horned Lark (*Eremphila alpestris*), Brown-headed Cowbird (*Molothrus ater*)). Only a few prairie endemic grassland species such as Sprague’s Pipit (*Anthus spragueii*), Chestnut-collared Longspur and Baird’s Sparrow (*Ammodramus bairdii*) were among the most abundant in natural landscapes but largely absent in extensive agricultural landscapes.

### Conservation Implications

Our results highlight how a greater similarity between agriculture and the natural ecosystem results in a higher peak proportion of agriculture at which species diversity and functional diversity begin to decline. This is likely due in part to a larger community of species that tolerate the change in the landscape when crops and pasture are grown in an ecosystem with structurally similar natural and agricultural land covers, regardless of whether they actually use the agricultural land covers. The prairie region has more extensive agriculture than the eastern forest region with agriculture representing approximately 62% of the land cover in the former and 32% in the latter. This difference in extent, combined with the differing contrast between the agricultural and natural land covers in the two ecosystems, points to different recommendations for the management of biodiversity in each case (Cunningham et al. 2013). When the agricultural extent is low but the structural contrast is high (i.e. Eastern Hardwood and Boreal Forest region) management practices should limit the expansion of high contrast agriculture, maintain existing semi-natural habitats (e.g. hedge rows) and restrict other human land uses in the broader landscape to the extent possible. When agriculture has a wide extent but low structural contrast with the native ecosystem (i.e. Prairie Pothole region) it requires practices that do not endanger ecosystem services, avoid overexploitation of the natural resource base and manage for native species intolerant of the production system (Cunningham et al. 2013). We emphasize this latter point in particular because prairie grasslands are among the most threatened ecosystems in North America (With et al. 2008, Grand et al. 2019). While a number of species in the prairie community tolerate some agriculture, several endemic species sensitive to agriculture are in serious decline with some such as Chestnut-collared Longspur and Lark Bunting (*Calamospiza melanocorys*) declining by more than 80% since the 1970s (Wilson et al. 2018). Other non-avian prairie endemic taxa intolerant to agriculture face similar threats and are federally listed as threatened or endangered (Environment and Climate Change Canada 2017). Conversely, some of the most threatened avian species in naturally forested ecosystems in eastern North America are also grassland species such as Bobolink (*Dolichonyx oryzivorus*) and Eastern Meadowlark (*Sturnella magna*) where reforestation following earlier forest loss for agriculture poses a risk of habitat loss. These examples highlight how in both ecosystems, managing for higher diversity will not necessarily require the same strategies or a focus on the same areas as managing for species in decline (Norris and Harper 2004, Wilson et al. 2019). A valuable area of future study in both ecosystems would be to identify what composition and configurations of agricultural and natural land covers allow us to best optimize the conservation potential for both total diversity and threatened species. It would also be valuable to examine how the relationships we studied are influenced by the history of agriculture in a region (Martin et al. 2012). The expansion of agriculture in the western hemisphere is relatively recent compared to the eastern hemisphere, particularly Europe and Asia. We hypothesize that the influence of similarity between natural and agricultural land covers on biodiversity may lessen with greater history of agriculture as some species adapt while those unable to eventually disappear.

## DATA DEPOSITION STATEMENT

Our work will be submitted to the Dryad Data Repository upon acceptance.

## Notes

### Competing Interest Statement

The authors have declared no competing interest.

