## Supplemental Material for "Extent of similarity between agricultural and natural land covers shapes how biodiversity responds to agricultural expansion at landscape scales"

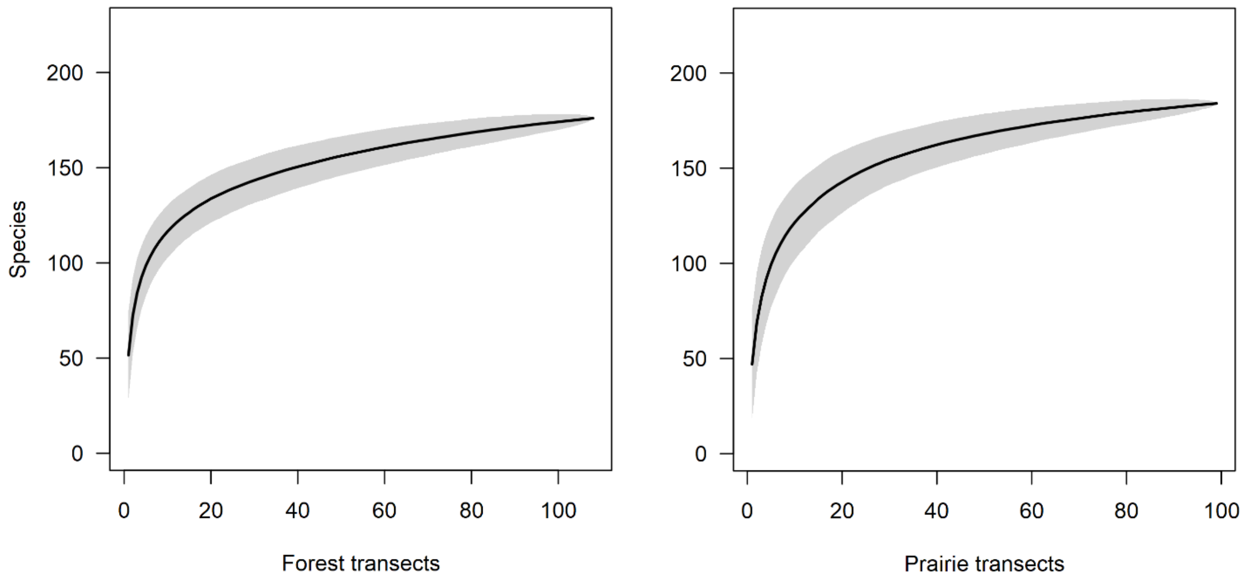


Figure S1. Species accumulation curve showing the number of species detected (with 95% uncertainty in light gray) with the random selection of any of up to 108 transects in the Forest ecosystem and 99 transects in the Prairie ecosystem.

Table S1. Mean percent of agricultural and natural land covers in 20km$^2$ landscapes surrounding transects in the prairie (n=203 landscape) and forest (n=220) ecosystems in this study. Percentages below indicate the approximate representation of cover types in each ecosystem. For analysis we randomly selected a subset of these transects in each ecoystem for a balanced comparison of how species diversity, functional diversity and abundance respond to an increasing extent of agriculture in each ecosystem. See Table S2 for individual crop types in the arable crops class.

| Land Cover | Prairie | Forest |
|---|---|---|
| Arable crops | 50.6 | 15.32 |
| Grassland | 22.4 | 0.08 |
| Pasture | 11.3 | 15.59 |
| Wetland | 3.3 | 3.74 |
| Broadleaf forest | 3.3 | 16.28 |
| Urban | 2.9 | 7.01 |
| Shrubland | 2.7 | 9.68 |
| Barren | 2.0 | 0.98 |
| Water | 1.3 | 3.12 |
| Coniferous forest | 0.3 | 5.58 |
| Mixed forest | 0.0 | 22.61 |

Table S2. Mean percent of total land cover and total agricultural cover represented by specific crop types in the prairie (n=203 landscape) and forest (n=220) ecosystems in this study.

| | Prairie | | Forest | |
|---|---|---|---|---|
| Land cover | % Total | % Agriculture | % Total | % Agriculture |
| Wheat | 27.21 | 37.63 | 0.21 | 0.64 |
| Pasture | 17.26 | 29.17 | 16.52 | 52.37 |
| Canola | 9.89 | 12.78 | 0.03 | 0.10 |
| Barley | 4.48 | 6.36 | 0.10 | 0.32 |
| Fallow | 2.58 | 3.83 | 0.00 | 0.01 |
| Peas | 2.25 | 2.70 | 0.00 | 0.00 |
| Lentils | 1.74 | 1.89 | 0.00 | 0.00 |
| Flaxseed | 1.22 | 1.48 | 0.00 | 0.00 |
| Winter Wheat | 0.86 | 1.54 | 0.00 | 0.00 |
| Mustard | 0.39 | 0.51 | 0.00 | 0.00 |
| Oats | 0.29 | 0.43 | 0.28 | 0.86 |
| Corn | 0.20 | 0.28 | 7.41 | 23.61 |

| | Prairie | | Forest | |
|---|---|---|---|---|
| Land cover | % | % | % | % |
| | Total | Agriculture | Total | Agriculture |
| Sunflower | 0.18 | 0.25 | 0.00 | 0.00 |
| Beans | 0.16 | 0.22 | 0.13 | 0.41 |
| Potatoes | 0.14 | 0.18 | 0.02 | 0.06 |
| Ray | 0.13 | 0.18 | 0.00 | 0.00 |
| Sugarbeets | 0.13 | 0.17 | 0.00 | 0.00 |
| Soybean | 0.11 | 0.17 | 5.00 | 15.90 |
| Canaryseed | 0.06 | 0.06 | 0.00 | 0.00 |
| Hemp | 0.03 | 0.04 | 0.00 | 0.02 |
| Tricale | 0.02 | 0.02 | 0.00 | 0.00 |
| Millet | 0.01 | 0.01 | 0.00 | 0.00 |
| Undifferentiated cereals | 0.00 | 0.00 | 1.46 | 4.66 |
| Vegetables | 0.00 | 0.00 | 0.18 | 0.57 |
| Undifferentiated crops | 0.00 | 0.00 | 0.06 | 0.19 |
| Orchards | 0.00 | 0.00 | 0.06 | 0.19 |

| | Prairie | | Forest | |
|---|---|---|---|---|
| Land cover | % Total | % Agriculture | % Total | % Agriculture |
| Nursery | 0.00 | 0.00 | 0.01 | 0.03 |
| Berries | 0.00 | 0.00 | 0.01 | 0.03 |
| Fruits | 0.00 | 0.00 | 0.00 | 0.02 |
| Buckwheat | 0.00 | 0.00 | 0.00 | 0.01 |

Table S3. Species included in the analysis along with their size (mass in grams), migratory, dietary and foraging strata guilds, and mean number detected per transect (based on 108 transects in the Forest ecosystem and 99 in the Prairie ecosystem). Migratory guilds (Mig) include N = Neotropical migrant, S = short-distance migrant, R = resident. For species with two migratory guild designations, the first refers to the forest and the second the prairie. Dietary guilds include C = Carnivore, I = Insectivore, N = Nectarivore, F = Frugivore, G = Granivore, H = Herbivore, O = Omnivore. Foraging strata guilds include A = Aerial Forager, B = Bark forager, G = Ground forager, LF = Lower foliage gleaner, SW = Surface water forager, UF = Upper foliage gleaner, UW = Underwater forager. Habitat guilds included A = Aquatic, T = Terrestrial. Nesting Guilds include G=Open cup on ground <0.5m, O = Open cup off ground (>0.5m), TC = Tree Cavity, RC = Rock cavity/Cliffs, BP = Brood Parasite. Species that primarily nest in association with human structures today were assigned based on their natural associations.

| **Species name (guild)** | **Scientific name** | **Size (g)** | **Mig** | **Diet** | **Strata** | **Habitat** | **Nest** | **Mean Forest** | **Mean Prairie** |
|---|---|---|---|---|---|---|---|---|---|
| American Black Duck | *Anas rubripes* | 1200 | S | H | SW | A | G | 0.11 | 0.00 |
| Alder Flycatcher | *Empidonax alnorum* | 13.5 | N | I | LF | T | O | 1.35 | 0.18 |
| American Avocet | *Recurvirostra americana* | 315 | S | I | SW | A | G | 0.00 | 0.35 |

| Species name (guild) | Scientific name | Size (g) | Mig | Diet | Strata | Habitat | Nest | Mean Forest | Mean Prairie |
|---|---|---|---|---|---|---|---|---|---|
| American Bittern | *Botaurus lentiginosus* | 700 | S | C | SW | A | G | 0.41 | 0.53 |
| American Coot | *Fulica americana* | 650 | S | H | SW | A | G | 0.00 | 1.99 |
| American Crow | *Corvus brachyrhynchos* | 450 | R/S | O | G | T | O | 13.74 | 5.25 |
| American Goldfinch | *Spinus tristis* | 13 | R/S | G | LF | T | O | 3.31 | 1.54 |
| American Kestrel | *Falco sparverius* | 117 | S | I | G | T | TC | 0.12 | 0.12 |
| American Redstart | *Setophaga ruticilla* | 8.3 | N | I | UF | T | O | 0.90 | 0.19 |
| American Robin | *Turdus migratorius* | 77 | S | I | LF | T | O | 14.02 | 3.70 |
| American Wigeon | *Mareca americana* | 720 | S | H | SW | A | G | 0.00 | 0.52 |
| American Woodcock | *Scolopax minor* | 200 | S | I | G | T | G | 0.04 | 0.00 |
| American White Pelican | *Pelecanus erythrorhynchos* | 7500 | S | C | UW | A | G | 0.00 | 0.45 |

| Species name (guild) | Scientific name | Size (g) | Mig | Diet | Strata | Habitat | Nest | Mean Forest | Mean Prairie |
|---|---|---|---|---|---|---|---|---|---|
| Bald Eagle | *Haliaeetus leucocephalus* | 4325 | S | C | SW | A | O | 0.02 | 0.02 |
| Bank Swallow | *Riparia riparia* | 13.5 | N | I | A | T | RC | 0.48 | 0.96 |
| Baltimore Oriole | *Icterus galbula* | 33 | N | I | UF | T | O | 0.72 | 0.66 |
| Barred Owl | *Strix varia* | 720 | R | C | G | T | TC | 0.01 | 0.00 |
| Barn Swallow | *Hirundo rustica* | 19 | N | I | A | T | RC | 2.32 | 2.02 |
| Baird's Sparrow | *Ammodramus bairdii* | 17.5 | S | G | G | T | G | 0.00 | 1.46 |
| Black-and-white Warbler | *Mniotilta varia* | 10.7 | N | I | B | T | G | 0.73 | 0.06 |
| Black-billed Cuckoo | *Coccyzus erythropthalmus* | 52 | N | I | UF | T | O | 0.20 | 0.26 |
| Black-billed Magpie | *Pica hudsonia* | 175 | R | O | G | T | O | 0.00 | 3.06 |
| Bay-breasted Warbler | *Setophaga castanea* | 12.5 | N | I | UF | T | O | 0.00 | 0.00 |

| Species name (guild) | Scientific name | Size (g) | Mig | Diet | Strata | Habitat | Nest | Mean Forest | Mean Prairie |
|---|---|---|---|---|---|---|---|---|---|
| Black-capped Chickadee | *Poecile atricapillus* | 11 | R | I | UF | T | TC | 3.09 | 0.31 |
| Black-crowned Night-Heron | *Nycticorax nycticorax* | 870 | S | C | SW | A | O | 0.05 | 0.04 |
| Belted Kingfisher | *Megaceryle alcyon* | 150 | S | C | SW | A | RC | 0.24 | 0.04 |
| Blue-gray Gnatcatcher | *Cinclidium frontale* | 6 | N | I | LF | T | O | 0.00 | 0.00 |
| Brown-headed Cowbird | *Molothrus ater* | 44 | S | G | G | T | BP | 1.08 | 7.66 |
| Blue-headed Vireo | *Vireo solitarius* | 16 | N | I | UF | T | O | 0.19 | 0.00 |
| Blackburnian Warbler | *Setophaga fusca* | 9.8 | N | I | UF | T | O | 0.27 | 0.00 |
| Blue Jay | *Cyanocitta cristata* | 85 | R | O | G | T | O | 1.90 | 0.08 |
| Black Tern | *Chlidonias niger* | 62 | N | I | SW | A | G | 0.00 | 1.04 |

| **Species name (guild)** | **Scientific name** | **Size (g)** | **Mig** | **Diet** | **Strata** | **Habitat** | **Nest** | **Mean Forest** | **Mean Prairie** |
|---|---|---|---|---|---|---|---|---|---|
| Black-necked Stilt | *Himantopus mexicanus* | 160 | N | I | SW | A | G | 0.00 | 0.04 |
| Bobolink | *Dolichonyx oryzivorus* | 43 | N | G | G | T | G | 3.24 | 1.09 |
| Boreal Chickadee | *Poecile hudsonicus* | 10 | R | I | UF | T | TC | 0.01 | 0.00 |
| Brewer's Blackbird | *Euphagus cyanocephalus* | 63 | S | I | G | T | O | 0.06 | 6.81 |
| Brown Creeper | *Certhia americana* | 8.4 | R | I | B | T | O | 0.04 | 0.00 |
| Brewer's Sparrow | *Spizella breweri* | 10.5 | S | G | G | T | O | 0.00 | 0.09 |
| Brown Thrasher | *Toxostoma rufum* | 69 | S | I | G | T | O | 0.49 | 0.36 |
| Black-throated Blue Warbler | *Setophaga caerulescens* | 10.2 | N | I | UF | T | O | 0.20 | 0.00 |
| Black-throated Green Warbler | *Setophaga virens* | 8.8 | N | I | UF | T | O | 0.60 | 0.00 |

| **Species name (guild)** | **Scientific name** | **Size (g)** | **Mig** | **Diet** | **Strata** | **Habitat** | **Nest** | **Mean Forest** | **Mean Prairie** |
|---|---|---|---|---|---|---|---|---|---|
| Bufflehead | *Bucephala albeola* | 380 | S | I | UW | A | TC | 0.00 | 0.27 |
| Bullock's Oriole | *Icterus bullockii* | 36 | N | I | UF | T | O | 0.00 | 0.02 |
| Broad-winged Hawk | *Buteo platypterus* | 390 | N | C | G | T | O | 0.05 | 0.00 |
| Blue-winged Teal | *Spatula discors* | 380 | N | H | SW | A | G | 0.02 | 2.72 |
| Blue-winged Warbler | *Vermivora cyanoptera* | 8.5 | N | I | LF | T | O | 0.01 | 0.00 |
| Canada Goose | *Branta canadensis* | 3050 | S | H | G | A | G | 7.17 | 5.90 |
| California Gull | *Larus californicus* | 610 | S | O | G | A | G | 0.00 | 0.41 |
| Canvasback | *Aythya valisineria* | 1220 | S | H | UW | A | G | 0.00 | 0.50 |
| Caspian Tern | *Hydroprogne caspia* | 660 | S | C | UW | A | G | 0.03 | 0.00 |
| Canada Warbler | *Cardellina canadensis* | 10.3 | N | I | LF | T | G | 0.07 | 0.00 |

| **Species name (guild)** | **Scientific name** | **Size (g)** | **Mig** | **Diet** | **Strata** | **Habitat** | **Nest** | **Mean Forest** | **Mean Prairie** |
|---|---|---|---|---|---|---|---|---|---|
| Chestnut-collared Longspur | *Calcarius ornatus* | 19 | S | G | G | T | G | 0.00 | 0.89 |
| Clay-colored Sparrow | *Spizella pallida* | 12 | S | G | G | T | G | 0.07 | 8.25 |
| Cedar Waxwing | *Bombycilla cedrorum* | 32 | S | F | UF | T | O | 2.51 | 1.19 |
| Cerulean Warbler | *Setophaga cerulea* | 9.3 | N | I | UF | T | O | 0.00 | 0.00 |
| Chipping Sparrow | *Spizella passerina* | 12 | S | G | G | T | O | 4.47 | 0.50 |
| Chimney Swift | *Chaetura pelagica* | 23 | N | I | A | T | TC | 0.02 | 0.00 |
| Cinnamon Teal | *Spatula cyanoptera* | 400 | S | H | SW | A | G | 0.00 | 0.03 |
| Cliff Swallow | *Petrochelidon pyrrhonota* | 21 | N | I | A | T | RC | 0.38 | 4.81 |
| Cape May Warbler | *Setophaga tigrina* | 11 | N | I | UF | T | O | 0.04 | 0.00 |
| Common Goldeneye | *Bucephala clangula* | 850 | R | I | UW | A | TC | 0.02 | 0.14 |

| Species name (guild) | Scientific name | Size (g) | Mig | Diet | Strata | Habitat | Nest | Mean Forest | Mean Prairie |
|---|---|---|---|---|---|---|---|---|---|
| Common Grackle | *Quiscalus quiscula* | 115 | S | O | G | T | O | 7.51 | 2.39 |
| Cooper's Hawk | *Accipiter cooperii* | 450 | S | C | G | T | O | 0.02 | 0.00 |
| Common Loon | *Gavia immer* | 4100 | S | C | UW | A | G | 0.18 | 0.02 |
| Common Merganser | *Mergus merganser* | 1530 | S | C | UW | A | G | 0.11 | 0.01 |
| Common Nighthawk | *Chordeiles minor* | 62 | N | I | A | T | G | 0.02 | 0.17 |
| Common Raven | *Corvus corax* | 1200 | R | O | G | T | RC | 0.81 | 1.37 |
| Common Tern | *Sterna hirundo* | 120 | N | C | UW | A | G | 0.00 | 0.05 |
| Common Yellowthroat | *Geothlypis trichas* | 10 | N | I | LF | T | G | 4.32 | 0.92 |
| Chestnut-sided Warbler | *Setophaga pensylvanica* | 9.6 | N | I | UF | T | O | 1.53 | 0.01 |
| Double-crested Cormorant | *Phalacrocorax auritus* | 1700 | S | C | UW | A | G | 0.28 | 0.52 |

| Species name (guild) | Scientific name | Size (g) | Mig | Diet | Strata | Habitat | Nest | Mean Forest | Mean Prairie |
|---|---|---|---|---|---|---|---|---|---|
| Dark-eyed Junco | *Junco hyemalis* | 19 | S | G | G | T | G | 0.09 | 0.03 |
| Downy Woodpecker | *Dryobates pubescens* | 27 | R | I | A | T | TC | 0.36 | 0.12 |
| Dusky Flycatcher | *Empidonax oberholseri* | 10.3 | N | I | LF | T | O | 0.00 | 0.04 |
| Eastern Bluebird | *Sialia sialis* | 31 | S | I | G | T | TC | 0.19 | 0.10 |
| Eared Grebe | *Podiceps nigricollis* | 300 | S | I | UW | A | G | 0.00 | 0.86 |
| Eastern Kingbird | *Tyrannus tyrannus* | 40 | N | I | A | T | O | 1.12 | 1.35 |
| Eastern Meadowlark | *Sturnella magna* | 90 | S | I | G | T | G | 1.79 | 0.00 |
| Eastern Phoebe | *Sayornis phoebe* | 20 | S | I | LF | T | RC | 0.88 | 0.13 |
| Eastern Towhee | *Pipilo erythrophthalmus* | 40 | S | G | G | T | G | 0.18 | 0.05 |
| Eurasian Collared Dove | *Streptopelia decaocto* | 200 | R | G | G | T | O | 0.01 | 0.07 |

| Species name (guild) | Scientific name | Size (g) | Mig | Diet | Strata | Habitat | Nest | Mean Forest | Mean Prairie |
|---|---|---|---|---|---|---|---|---|---|
| Eastern Screech Owl | *Megascops asio* | 180 | R | C | G | T | TC | 0.01 | 0.00 |
| European Starling | *Sturnus vulgaris* | 82 | R | O | G | T | TC | 13.08 | 4.23 |
| Evening Grosbeak | *Hesperiphona vespertina* | 60 | R | G | UF | T | O | 0.01 | 0.00 |
| Eastern Wood Pewee | *Contopus virens* | 14 | N | I | A | T | O | 0.50 | 0.05 |
| Ferruginous Hawk | *Buteo regalis* | 1600 | S | C | G | T | O | 0.00 | 0.09 |
| Field Sparrow | *Spizella pusilla* | 12.5 | S | G | G | T | G | 0.31 | 0.00 |
| Fox Sparrow | *Passerella iliaca* | 32 | S | G | G | T | G | 0.01 | 0.00 |
| Forster's Tern | *Sterna forsteri* | 160 | S | I | UW | A | G | 0.00 | 0.03 |
| Franklin's Gull | *Leucophaeus pipixcan* | 280 | N | O | G | A | G | 0.00 | 5.69 |
| Gadwall | *Mareca strepera* | 910 | S | H | SW | A | G | 0.04 | 1.71 |

| **Species name (guild)** | **Scientific name** | **Size (g)** | **Mig** | **Diet** | **Strata** | **Habitat** | **Nest** | **Mean Forest** | **Mean Prairie** |
|---|---|---|---|---|---|---|---|---|---|
| Great Black-backed Gull | *Larus marinus* | 1650 | S | O | G | A | G | 0.02 | 0.00 |
| Great Blue Heron | *Ardea herodias* | 2400 | S | C | SW | A | O | 0.35 | 0.16 |
| Great Crested Flycatcher | *Myiarchus crinitus* | 34 | N | I | UF | T | TC | 0.88 | 0.14 |
| Golden-crowned Kinglet | *Regulus satrapa* | 6 | S | I | UF | T | O | 0.10 | 0.00 |
| Great Gray Owl | *Strix nebulosa* | 1080 | R | C | G | T | G | 0.00 | 0.01 |
| Great Horned Owl | *Bubo virginianus* | 1400 | R | C | G | T | O | 0.01 | 0.24 |
| Golden Eagle | *Aquila chrysaetos* | 4575 | S | C | G | T | RC | 0.00 | 0.01 |
| Gray Catbird | *Dumetella carolinensis* | 37 | N | I | LF | T | O | 0.69 | 0.52 |
| Great Egret | *Ardea alba* | 870 | S | C | SW | A | O | 0.01 | 0.00 |
| Green Heron | *Butorides virescens* | 210 | S | C | SW | A | O | 0.19 | 0.00 |

| Species name (guild) | Scientific name | Size (g) | Mig | Diet | Strata | Habitat | Nest | Mean Forest | Mean Prairie |
|---|---|---|---|---|---|---|---|---|---|
| Gray Jay | *Perisoreus canadensis* | 70 | R | O | G | T | O | 0.03 | 0.00 |
| Gray Partridge | *Perdix perdix* | 390 | R | H | G | T | G | 0.01 | 0.16 |
| Grasshopper Sparrow | *Ammodramus savannarum* | 17 | S | I | G | T | G | 0.10 | 0.47 |
| Green-winged Teal | *Anas carolinensis* | 350 | S | H | SW | A | G | 0.00 | 0.54 |
| Golden-winged Warbler | *Vermivora chrysoptera* | 8.8 | N | I | LF | T | G | 0.02 | 0.01 |
| Hairy Woodpecker | *Dryobates villosus* | 66 | R | I | B | T | TC | 0.28 | 0.07 |
| Herring Gull | *Larus smithsonianus* | 1150 | S | C | SW | A | G | 1.61 | 0.00 |
| Hermit Thrush | *Catharus guttatus* | 31 | S | I | G | T | G | 0.85 | 0.03 |
| House Finch | *Haemorhous mexicanus* | 21 | R | G | LF | T | O | 0.24 | 0.01 |
| Horned Grebe | *Podiceps auritus* | 450 | S | C | UW | A | G | 0.00 | 0.25 |

| Species name (guild) | Scientific name | Size (g) | Mig | Diet | Strata | Habitat | Nest | Mean Forest | Mean Prairie |
|---|---|---|---|---|---|---|---|---|---|
| Horned Lark | *Eremophila alpestris* | 32 | S | G | G | T | G | 0.74 | 4.98 |
| Hooded Merganser | *Lophodytes cucullatus* | 620 | S | O | UW | A | TC | 0.09 | 0.05 |
| House Sparrow | *Passer domesticus* | 28 | R | I | G | T | TC | 2.74 | 6.46 |
| House Wren | *Troglodytes aedon* | 11 | N | I | LF | T | TC | 1.07 | 3.47 |
| Indigo Bunting | *Passerina cyanea* | 14.5 | N | G | LF | T | O | 0.92 | 0.02 |
| Killdeer | *Charadrius vociferus* | 95 | S | I | G | T | G | 1.26 | 1.85 |
| King Rail | *Rallus elegans* | 360 | S | O | SW | A | G | 0.01 | 0.00 |
| Lark Bunting | *Calamospiza melanocorys* | 38 | S | I | G | T | G | 0.00 | 1.43 |
| Lark Sparrow | *Chondestes grammacus* | 29 | S | G | G | T | G | 0.00 | 0.13 |
| Long-billed Curlew | *Numenius americanus* | 590 | S | I | G | T | G | 0.00 | 0.38 |

| **Species name (guild)** | **Scientific name** | **Size (g)** | **Mig** | **Diet** | **Strata** | **Habitat** | **Nest** | **Mean Forest** | **Mean Prairie** |
|---|---|---|---|---|---|---|---|---|---|
| Le Conte's Sparrow | *Ammodramus leconteii* | 13 | S | G | G | T | G | 0.00 | 0.43 |
| Least Bittern | *Ixobrychus exilis* | 80 | S | C | SW | A | G | 0.01 | 0.00 |
| Least Flycatcher | *Empidonax minimus* | 10.3 | N | I | A | T | O | 0.59 | 1.55 |
| Lesser Scaup | *Aythya affinis* | 830 | S | H | UW | A | G | 0.00 | 0.97 |
| Lesser Yellowlegs | *Tringa flavipes* | 80 | N | I | SW | A | G | 0.00 | 0.13 |
| Lincoln's Sparrow | *Melospiza lincolnii* | 17 | S | G | G | T | G | 0.03 | 0.03 |
| Loggerhead Shrike | *Lanius ludovicianus* | 48 | S | I | G | T | O | 0.00 | 0.20 |
| Marbled Godwit | *Limosa fedoa* | 370 | S | I | G | A | G | 0.00 | 1.07 |
| Mallard | *Anas platyrhynchos* | 1100 | S | O | SW | A | G | 1.63 | 6.04 |
| Magnolia Warbler | *Setophaga magnolia* | 8.7 | N | I | UF | T | O | 0.50 | 0.02 |

| **Species name (guild)** | **Scientific name** | **Size (g)** | **Mig** | **Diet** | **Strata** | **Habitat** | **Nest** | **Mean Forest** | **Mean Prairie** |
|---|---|---|---|---|---|---|---|---|---|
| Marsh Wren | *Cistothorus palustris* | 11 | S | I | LF | A | O | 0.07 | 0.45 |
| McCown's Longspur | *Rhynchophanes mccownii* | 23 | S | G | G | T | G | 0.00 | 0.04 |
| Merlin | *Falco columbarius* | 190 | S/R | C | A | T | RC | 0.01 | 0.08 |
| Mountain Bluebird | *Sialia currucoides* | 29 | S | I | G | T | TC | 0.00 | 0.13 |
| Mourning Dove | *Zenaida macroura* | 120 | S | G | G | T | O | 4.68 | 4.10 |
| Mourning Warbler | *Geothlypis philadelphia* | 12.5 | N | I | LF | T | G | 0.38 | 0.01 |
| Myrtle Warbler | *Setophaga coronata* | 12.3 | S | I | UF | T | O | 0.58 | 0.02 |
| Nashville Warbler | *Leiothlypis ruficapilla* | 8.7 | N | I | UF | T | G | 0.77 | 0.01 |
| Nelson's Sparrow | *Ammodramus nelsoni* | 17 | S | G | G | A | G | 0.00 | 0.15 |
| Northern Cardinal | *Cardinalis cardinalis* | 45 | R | G | G | T | O | 0.93 | 0.00 |

| **Species name (guild)** | **Scientific name** | **Size (g)** | **Mig** | **Diet** | **Strata** | **Habitat** | **Nest** | **Mean Forest** | **Mean Prairie** |
|---|---|---|---|---|---|---|---|---|---|
| Northern Flicker | *Colaptes auratus* | 130 | S | I | G | T | TC | 0.61 | 0.30 |
| Northern Goshawk | *Accipiter gentilis* | 950 | R | C | G | T | O | 0.02 | 0.00 |
| Northern Harrier | *Circus hudsonius* | 420 | S | C | G | T | G | 0.09 | 0.35 |
| Northern Mockingbird | *Mimus polyglottos* | 49 | S | I | G | T | O | 0.01 | 0.00 |
| Northern Parula | *Setophaga americana* | 8.6 | N | I | UF | T | O | 0.08 | 0.00 |
| Northern Pintail | *Anas acuta* | 800 | S | H | SW | A | G | 0.00 | 1.34 |
| Northern Shoveler | *Spatula clypeata* | 610 | S | I | SW | A | G | 0.00 | 1.48 |
| Northern Waterthrush | *Parkesia noveboracensis* | 19 | N | I | G | A | G | 0.23 | 0.03 |
| Northern Rough-winged Swallow | *Stelgidopteryx serripennis* | 16 | N | I | A | T | RC | 0.18 | 0.00 |

| **Species name (guild)** | **Scientific name** | **Size (g)** | **Mig** | **Diet** | **Strata** | **Habitat** | **Nest** | **Mean Forest** | **Mean Prairie** |
|---|---|---|---|---|---|---|---|---|---|
| Northern Saw-whet Owl | *Aegolius acadicus* | 80 | S | C | G | T | TC | 0.01 | 0.00 |
| Orange-crowned Warbler | *Leiothlypis celata* | 9 | N | I | UF | T | G | 0.00 | 0.03 |
| Orchard Oriole | *Icterus spurius* | 19 | N | F | LF | T | O | 0.01 | 0.06 |
| Olive-sided Flycatcher | *Contopus cooperi* | 32 | N | I | A | T | O | 0.02 | 0.00 |
| Osprey | *Pandion haliaetus* | 1600 | S | C | SW | A | O | 0.06 | 0.00 |
| Ovenbird | *Seiurus aurocapilla* | 19.5 | N | I | G | T | G | 2.73 | 0.07 |
| Pied-billed Grebe | *Podilymbus podiceps* | 450 | S | I | UW | A | G | 0.03 | 0.46 |
| Philadelphia Vireo | *Vireo philadelphicus* | 12 | N | I | UF | T | O | 0.05 | 0.04 |
| Pine Siskin | *Spinus pinus* | 15 | R | G | UF | T | O | 0.20 | 0.03 |
| Pine Warbler | *Setophaga pinus* | 12 | S | I | UF | T | O | 0.33 | 0.00 |

| **Species name (guild)** | **Scientific name** | **Size (g)** | **Mig** | **Diet** | **Strata** | **Habitat** | **Nest** | **Mean Forest** | **Mean Prairie** |
|---|---|---|---|---|---|---|---|---|---|
| Pileated Woodpecker | *Dryocopus pileatus* | 290 | R | I | B | T | TC | 0.24 | 0.05 |
| Prairie Falcon | *Falco mexicanus* | 720 | R | C | G | T | RC | 0.00 | 0.01 |
| Purple Finch | *Haemorhous purpureus* | 25 | S | G | UF | T | O | 0.29 | 0.01 |
| Purple Martin | *Progne subis* | 56 | N | I | A | T | TC | 0.15 | 0.30 |
| Rose-breasted Grosbeak | *Pheucticus ludovicianus* | 45 | N | I | UF | T | O | 0.76 | 0.19 |
| Ring-billed Gull | *Larus delawarensis* | 520 | S | O | G | A | G | 7.46 | 1.83 |
| Red-breasted Merganser | *Mergus serrator* | 1060 | S | C | UW | A | G | 0.01 | 0.00 |
| Red-breasted Nuthatch | *Sitta canadensis* | 10 | R | I | B | T | TC | 0.46 | 0.02 |
| Red-bellied Woodpecker | *Melanerpes carolinus* | 63 | R | O | B | T | TC | 0.09 | 0.00 |
| Ruby-crowned Kinglet | *Regulus calendula* | 6.5 | N | I | UF | T | O | 0.12 | 0.00 |

| **Species name (guild)** | **Scientific name** | **Size (g)** | **Mig** | **Diet** | **Strata** | **Habitat** | **Nest** | **Mean Forest** | **Mean Prairie** |
|---|---|---|---|---|---|---|---|---|---|
| Red Crossbill | *Loxia curvirostra* | 36 | R | G | UF | T | O | 0.00 | 0.02 |
| Redhead | *Aythya americana* | 1050 | S | H | UW | A | G | 0.00 | 0.68 |
| Red-eyed Vireo | *Vireo olivaceus* | 17 | N | I | UF | T | O | 4.89 | 0.63 |
| Red-headed Woodpecker | *Melanerpes erythrocephalus* | 72 | S | I | UF | T | TC | 0.01 | 0.04 |
| Ring-necked Duck | *Aythya collaris* | 700 | S | H | SW | A | G | 0.02 | 0.09 |
| Red-necked Grebe | *Podiceps grisegena* | 1000 | S | C | UW | A | G | 0.00 | 0.13 |
| Ring-necked Pheasant | *Scleroptila streptophora* | 1150 | R | G | G | T | G | 0.08 | 0.49 |
| Red-naped Sapsucker | *Sphyrapicus nuchalis* | 50 | S | I | B | T | TC | 0.00 | 0.01 |
| Rock Pigeon | *Columbia livia* | 270 | R | G | G | T | RC | 2.25 | 1.85 |

| Species name (guild) | Scientific name | Size (g) | Mig | Diet | Strata | Habitat | Nest | Mean Forest | Mean Prairie |
|---|---|---|---|---|---|---|---|---|---|
| Red-shouldered Hawk | *Buteo lineatus* | 630 | S | C | G | T | O | 0.01 | 0.00 |
| Red-tailed Hawk | *Buteo jamaicensis* | 1080 | S | C | G | T | O | 0.09 | 0.60 |
| Ruby-throated Hummingbird | *Archilochus colubris* | 3.2 | N | N | LF | T | O | 0.11 | 0.03 |
| Rusty Blackbird | *Euphagus carolinus* | 60 | S | I | G | T | O | 0.00 | 0.01 |
| Ruddy Duck | *Oxyura jamaicensis* | 560 | S | I | UW | A | G | 0.00 | 0.70 |
| Ruffed Grouse | *Bonasa umbellus* | 580 | R | H | G | T | G | 0.11 | 0.12 |
| Red-winged Blackbird | *Agelaius phoeniceus* | 52 | S | I | G | T | O | 16.18 | 24.96 |
| Sandhill Crane | *Antigone canadensis* | 4850 | S | O | G | A | G | 0.19 | 0.16 |
| Say's Phoebe | *Sayornis saya* | 21 | S | I | G | T | RC | 0.00 | 0.08 |

| **Species name (guild)** | **Scientific name** | **Size (g)** | **Mig** | **Diet** | **Strata** | **Habitat** | **Nest** | **Mean Forest** | **Mean Prairie** |
|---|---|---|---|---|---|---|---|---|---|
| Savannah Sparrow | *Passerculus sandwichensis* | 20 | S | G | G | T | G | 4.60 | 9.43 |
| Scarlet Tanager | *Piranga olivacea* | 28 | N | I | UF | T | O | 0.23 | 0.01 |
| Short-eared Owl | *Asio flammeus* | 350 | S | C | G | T | G | 0.00 | 0.09 |
| Sedge Wren | *Cistothorus stellaris* | 9 | S | I | LF | A | G | 0.01 | 0.48 |
| Sora | *Porzana carolina* | 75 | S | G | G | A | G | 0.04 | 1.27 |
| Song Sparrow | *Melospiza melodia* | 20 | S | G | G | T | G | 9.56 | 1.87 |
| Sprague's Pipit | *Anthus spragueii* | 25 | S | I | G | T | G | 0.00 | 1.47 |
| Spotted Sandpiper | *Actitis macularius* | 40 | N | I | G | A | G | 0.09 | 0.11 |
| Spotted Towhee | *Pipilo maculatus* | 40 | S | G | G | T | G | 0.00 | 0.04 |
| Sharp-shinned Hawk | *Accipiter striatus* | 140 | S | C | A | T | O | 0.07 | 0.03 |

| **Species name (guild)** | **Scientific name** | **Size (g)** | **Mig** | **Diet** | **Strata** | **Habitat** | **Nest** | **Mean Forest** | **Mean Prairie** |
|---|---|---|---|---|---|---|---|---|---|
| Sharp-tailed Grouse | *Tympanuchus phasianellus* | 880 | R | H | G | T | G | 0.00 | 0.43 |
| Swainson's Hawk | *Buteo swainsoni* | 855 | N | C | G | T | O | 0.00 | 0.61 |
| Swamp Sparrow | *Melospiza georgiana* | 17 | S | I | G | A | G | 0.60 | 0.02 |
| Swainson's Thrush | *Catharus ustulatus* | 31 | N | I | LF | T | O | 0.22 | 0.04 |
| Tennessee Warbler | *Oreothlypis peregrina* | 10 | N | I | HF | T | G | 0.04 | 0.02 |
| Tree Swallow | *Tachycineta bicolor* | 20 | S | I | A | T | TC | 1.48 | 1.47 |
| Tufted Titmouse | *Baeolophs bicolor* | 21.5 | R | I | HF | T | C | 0.01 | 0.00 |
| Turkey Vulture | *Cathartes aura* | 1830 | S | S | G | T | RC | 0.19 | 0.07 |
| Upland Sandpiper | *Bartramia longicauda* | 170 | N | I | G | T | G | 0.11 | 0.74 |
| Veery | *Catharus fuscescens* | 31 | N | I | G | T | G | 2.02 | 0.11 |

| Species name (guild) | Scientific name | Size (g) | Mig | Diet | Strata | Habitat | Nest | Mean Forest | Mean Prairie |
|---|---|---|---|---|---|---|---|---|---|
| Vesper Sparrow | *Pooecetes gramineus* | 26 | S | G | G | T | G | 0.28 | 6.77 |
| Virginia Rail | *Rallus limicola* | 85 | S | O | SW | A | G | 0.07 | 0.04 |
| Warbling Vireo | *Vireo gilvus* | 12 | N | I | UF | T | O | 1.40 | 1.36 |
| White-breasted Nuthatch | *Sitta carolinensis* | 21 | R | I | B | T | TC | 0.35 | 0.07 |
| White-crowned Sparrow | *Zonotrichia leucophrys* | 29 | S | G | G | T | G | 0.00 | 0.09 |
| Western Grebe | *Aechmophorus occidentalis* | 1500 | S | C | UW | A | G | 0.00 | 0.05 |
| Western Kingbird | *Tyrannus verticalis* | 40 | N | I | A | T | O | 0.00 | 0.68 |
| Western Meadowlark | *Sturnella neglecta* | 97 | S | I | G | T | G | 0.01 | 16.93 |
| Western Tanager | *Piranga ludoviciana* | 28 | N | I | UF | T | O | 0.00 | 0.01 |
| Willow Flycatcher | *Empidonax traillii* | 13.5 | N | I | A | T | O | 0.22 | 0.02 |

| **Species name (guild)** | **Scientific name** | **Size (g)** | **Mig** | **Diet** | **Strata** | **Habitat** | **Nest** | **Mean Forest** | **Mean Prairie** |
|---|---|---|---|---|---|---|---|---|---|
| Willet | *Tringa semipalmata* | 215 | N | I | G | A | G | 0.00 | 0.93 |
| Wilson's Phalarope | *Phalaropus tricolor* | 60 | N | I | SW | A | G | 0.00 | 1.25 |
| Wilson's Snipe | *Gallinago delicata* | 105 | S | I | G | A | G | 0.51 | 2.23 |
| Wild Turkey | *Meleagris gallopavo* | 5800 | R | H | G | T | G | 0.63 | 0.11 |
| Wilson's Warbler | *Cardellina pusilla* | 7.7 | N | I | LF | T | O | 0.01 | 0.00 |
| Winter Wren | *Troglodytes hiemalis* | 9 | S | I | G | T | G | 0.38 | 0.00 |
| Wood Duck | *Aix sponsa* | 600 | S | H | SW | A | TC | 0.20 | 0.14 |
| Wood Thrush | *Hylocichla mustelina* | 47 | N | I | G | T | O | 0.44 | 0.00 |
| White-throated Sparrow | *Zonotrichia albicollis* | 26 | S | G | G | T | G | 2.70 | 0.11 |
| White-winged Crossbill | *Loxia leucoptera* | 26 | R | G | UF | T | O | 0.02 | 0.01 |

| Species name (guild) | Scientific name | Size (g) | Mig | Diet | Strata | Habitat | Nest | Mean Forest | Mean Prairie |
|---|---|---|---|---|---|---|---|---|---|
| Western Wood Pewee | *Contopus sordidulus* | 13 | N | I | A | T | O | 0.00 | 0.07 |
| Yellow-billed Cuckoo | *Coccyzus americanus* | 65 | N | I | UF | T | O | 0.04 | 0.00 |
| Yellow-bellied Flycatcher | *Empidonax flaviventris* | 11.5 | N | I | UF | T | G | 0.01 | 0.00 |
| Yellow-bellied Sapsucker | *Sphyrapicus varius* | 50 | S | I | B | T | TC | 0.68 | 0.22 |
| Yellow-headed Blackbird | *Xanthocephalus xanthocephalus* | 65 | S | I | G | A | O | 0.00 | 3.83 |
| Yellow-throated Vireo | *Vireo flavifrons* | 18 | N | I | HF | T | O | 0.04 | 0.02 |
| Yellow Warbler | *Setophaga petechia* | 9.5 | N | I | LF | T | O | 1.97 | 2.76 |

Table S4. Ecosystem-specific means and coefficient of variation for species diversity (Shannon index), functional diversity (RaoQ) and abundance (individuals) on forest and prairie transects. Values after means are 95% confidence intervals.

| Measure | Forest | | Prairie | |
|---|---|---|---|---|
| | Mean | CV | Mean | CV |
| Species diversity | 3.21 (3.12 - 3.31) | 0.092 | 3.08 (2.96 - 3.20) | 0.118 |
| Functional diversity | 0.112 (0.106 - 0.118) | 0.163 | 0.101 (0.091 - 0.111) | 0.292 |
| Abundance | 188 (167 - 209) | 0.35 | 216 (188 - 245) | 0.39 |
